# Redundancy in synaptic connections enables neurons to learn optimally

**DOI:** 10.1101/127407

**Authors:** Naoki Hiratani, Tomoki Fukai

## Abstract

Recent experimental studies suggest that, in cortical microcircuits of the mammalian brain, the majority of neuron-to-neuron connections are realized by multiple synapses. However, it is not known whether such redundant synaptic connections provide any functional benefit. Here, we show that redundant synaptic connections enable near-optimal learning in cooperation with synaptic rewiring. By constructing a simple dendritic neuron model, we demonstrate that with multisynaptic connections, synaptic plasticity approximates a sample-based Bayesian filtering algorithm known as particle filtering, and wiring plasticity implements its resampling process. Applying the proposed framework to a detailed single neuron model, we show that the model accounts for many experimental observations, including the dendritic position dependence of spike-timing-dependent plasticity, and the functional synaptic organization on the dendritic tree based on the stimulus selectivity of presynaptic neurons. Our study provides a novel conceptual framework for synaptic plasticity and rewiring.

## Introduction

Synaptic connection between neurons is the fundamental substrate for learning and computation in neural circuits. Previous morphological studies suggest that in cortical microcircuits, often several synaptic connections are found between the presynaptic axons and the postsynaptic dendrites of two connected neurons (Deuchars et al., 1994; Markram et al., 1997; Feldmeyer et al., 1999). Recent connectomics studies confirmed these observations in somatosensory (Kasthuri et al., 2015), visual (Lee et al., 2016), and entorhinal (Schmidt et al., 2017) cortex, and also in hippocampus (Bartol et al., 2015). In particular, in barrel cortex, the average number of synapses per connection is estimated to be around 10 (Gal et al., 2017). However, the functional importance of multisynaptic connections remains unknown. Especially, from a computational perspective, such redundancy in connection structure is potentially harmful for learning due to degeneracy (Watanabe, 2001; Amari et al., 2006). In this work, we study how neurons perform learning with multisynaptic connections and whether redundancy provides any benefit, from a Bayesian perspective.

Bayesian framework has been established as a candidate principle of information processing in the brain (Knill and Pouget, 2004; Körding and Wolpert, 2006). Many results further suggest that not only computation, but learning process is also near optimal in terms of Bayesian for given stream of information (Behrens et al., 2007; Lake et al., 2015; Madarasz et al., 2016), yet its underlying plasticity mechanism remains largely elusive. Previous theoretical studies revealed that Hebbian-type plasticity rules eventually enable neural circuits to perform optimal computation under appropriate normalization (Soltani and Wang, 2010; Nessler et al., 2013). However, these rules are not optimal in terms of learning, so that the learning rates are typically too slow to perform learning from a limited number of observations. Recently, some learning rules are proposed for rapid learning (Aitchison and Latham, 2014; Gütig, 2016), yet their biological plausibility are still debatable. Here, we propose a novel framework of non-parametric near-optimal learning using multisynaptic connections. We show that neurons can exploit the variability among synapses in a multisynaptic connection to accurately estimate the causal relationship between pre and postsynaptic activity. The learning rule is first derived for a simple neuron model, and then implemented in a detailed single neuron model. The derived rule is consistent with many known properties of dendritic plasticity and synaptic organization, including a recent finding on the dendritic retinotopy in Layer 2/3 (L2/3) pyramidal neurons of rodent visual cortex (Iacaruso et al., 2017). Furthermore, the model reveals potential functional roles of anti-Hebbian synaptic plasticity observed in distal dendrites (Letzkus et al., 2006; Sjöström and Häusser, 2006).

## Results

### A conceptual model of learning with multisynaptic connections

Let us first consider a model of two neurons connected with *K* numbers of synapses (Fig. 1A) to illustrate the concept of the proposed framework. In the model, synaptic connections from the presynaptic neuron are distributed on the dendritic tree of the postsynaptic neuron as observed in experiments (Markram et al., 1997; Feldmeyer et al., 1999). Although a cortical neuron receives synaptic inputs from several thousands of presynaptic neurons in reality, here we consider the simplified model to illustrate the conceptual novelty of the proposed framework. More realistic models will be studied in following sections.

The synapses generate different amplitudes of excitatory postsynaptic potentials at the soma mainly through two mechanisms. First, the amplitude of dendritic attenuation varies from synapse to synapse, because the distances from the soma are different (Stuart and Spruston, 1998; Segev and London, 2000). Let us denote this dendritic position dependence of synapse *k* as *v*_*k*_, and call it as the unit EPSP, because *v*_*k*_ corresponds to the somatic potential caused by a unit conductance change at the synapse (i.e. somatic EPSP per AMPA receptor). As depicted in Figure 1A, unit EPSP *v*_*k*_ takes a small (large) value on a synapse at a distal (proximal) position on the dendrite. The second factor is the amount of AMPA receptors in the corresponding spine, which is approximately proportional to its spine size (Matsuzaki et al., 2004). If we denote this spine size factor as *g*_*k*_, the somatic EPSP caused by a synaptic input through synapse *k* is written as *w*_*k*_ = *g*_*k*_*v*_*k*_. This means that even if the synaptic contact is made at a distal dendrite (i.e. even if *v*_*k*_ is small), if the spine size *g*_*k*_ is large, a synaptic input through synapse *k* has a strong impact at the soma (e.g. red synapse in Fig. 1A) or vice versa (e.g. cyan synapse in Fig. 1A).

**Fig. 1.**
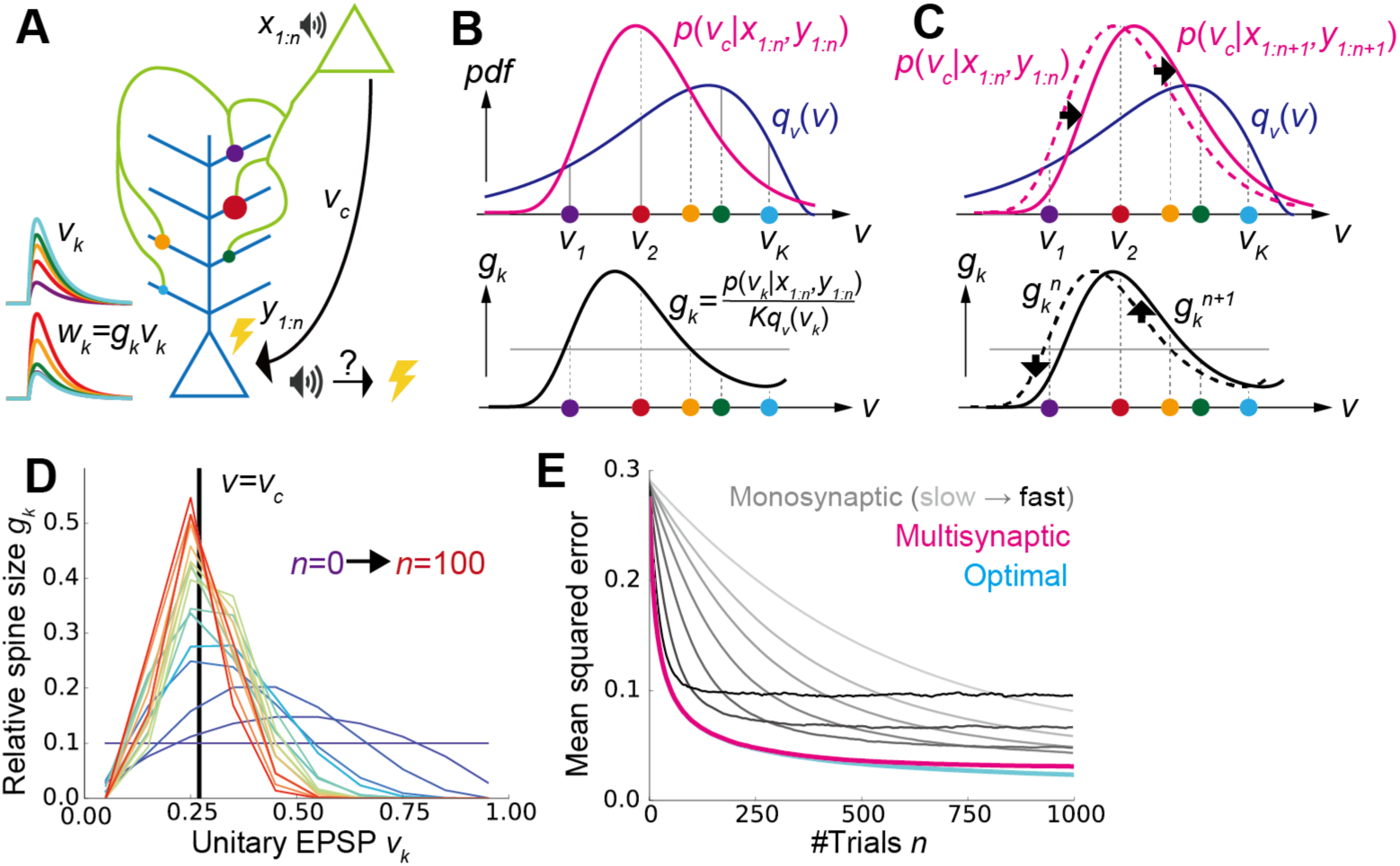
A conceptual model of multisynaptic learning. **A)** Schematic figure of the model consist of two neurons connected with *K* synapses. Curves on the left represent unit EPSP *v*_*k*_ (top) and the weighted EPSP *w*_*k*_=*g*_*k*_*v*_*k*_ (bottom) of each synaptic connection. Note that synapses are consistently colored throughout Figure 1 and 2. **B** Schematics of non-parametric representation of the probability distribution by multisynaptic connections. In both graphs, *x*-axes are unit EPSP, and the left (right) side corresponds to distal (proximal) dendrite. The mean over the true distribution *p*(*v*_*c*_|*x*_*1:n*_,*y*_*1:n*_) can be approximately calculated by taking samples (i.e. synapses) from the unit EPSP distribution *q*_*v*_(*v*) (top), and then taking a weighted sum over the spine size factor *g*_*k*_ representing the ratio *p*(*v*_*c*_|*x*_*1:n*_,*y*_*1:n*_)/*q*_*v*_(*v*) (bottom). **C)** Illustration of synaptic weight updating. When the distribution *p*(*v*_*c*_|*x*_*1:n+1*_,*y*_*1:n+1*_) comes to the right side of the original distribution *p*(*v*_*c*_|*x*_*1:n*_,*y*_*1:n*_), a synaptic weight *gk*^*n+1*^ become larger (smaller) than *g*_*k*_^*n*^ at proximal (distal) synapses. **D)** An example of learning dynamics at *K*=10 and *q*_*v*_(*v*)=const. Each curve represents the distribution of relative spine size *{*gk*}*, and the colors represent the growth of trial number. **E)** Comparison of performance among the proposed method, the monosynaptic rule, and the exact solution (see *A conceptual model of multisynaptic learning* in Methods for details). The monosynaptic learning rule was implemented with *η*=0.01, 0.015, 0.02, 0.03, 0.05, 0.1, 0.2 (from gray to black), and the initial value was taken as 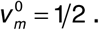. Lines were calculated by taking average over 10^4^ independent simulations.

On this model, we consider a simplified classical conditioning task as an example, though the framework is applicable for various inference tasks. Here, the presynaptic neuron activity represents the conditioned stimulus (CS) such as tone, and the postsynaptic neuron activity represents the unconditioned stimulus (US) such as shock. CS and US are represented by binary variables *x*_*n*_ ∈{0,1} and *y*_*n*_ ∈{0,1}, where *x*_*n*_ =1 (*y*_*n*_ =1) denotes the presence of the CS (US), and subscript *n* stands for the trial number (Fig. 1A). Learning behavior of animals and human in such a conditioning can be explained by the Bayesian framework (Courville et al., 2006). In particular, in order to invoke an appropriate behavioral response, the brain needs to keep track of the likelihood of US given CS *v*_*c*_ ≡ *p* (*y*_*n*_ = 1|*x*_*n*_ =1), presumably by changing the synaptic weight between corresponding neurons. Thus, we consider supervised learning of the conditional probability *v*_*c*_ by multisynaptic connections, from pre and postsynaptic activities representing US and CS, respectively. From finite trials up to *n*, this conditional probability is estimated as 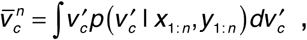, where *x*_*1:n*_={*x*_*1*_,*x*_*2*_,…,*x*_*n*_} and *y*_*1:n*_={*y*_*1*_,*y*_*2*_,…,*y*_*n*_} are the histories of input and output activities, and *p*(*v*_*c*_ | *x*_*1:n*_, *y*_*1:n*_) is the probability distribution of the hidden parameter *v*_*c*_ after *n* trials. Importantly, in general, it is impossible to get the optimal estimation of 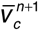 directly from 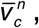 because in order to calculate 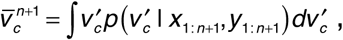 one first needs to calculate the distribution *p* (*v*_*c*_ | *x*_1:*n+1*_,*y*_1:*n+1*_) by integrating the previous distribution *p*(*v*_*c*_ | *x*_*1:n*_, *y*_*1:n*_) and the new observation at trial *n+1*: {*x*_*n+1*_, *y*_*n+1*_}. This means that for near-optimal learning, synaptic connections need to learn and represent the distribution *p*(*v*_*c*_ | *x*_*1:n*_,*y*_*1:n*_) instead of the point estimation 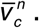. But, how can synapses achieve that? The key hypothesis of this paper is that redundancy in synaptic connections is the substrate for the non-parametric representation of this probabilistic distribution. Below, we show that dendritic summation over multisynaptic connections yields the optimal estimation from the given distribution *p*(*v*_*c*_ | *x*_*1:n*_, *y*_*1:n*_), and dendritic-position-dependent Hebbian synaptic plasticity updates this distribution.

### Dendritic summation as importance sampling

We first consider how dendritic summation achieves the calculation of the mean conditional probability 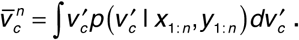. It is generally difficult to evaluate this integral by directly taking samples from the distribution *p*(*v*_*c*_ | *x*_*1:n*_,*y*_*1:n*_) in a biologically plausible way, because the cumulative distribution changes its shape at every trial. Nevertheless, we can still estimate the mean value by using an alternative distribution as the proposal distribution, and taking weighted samples from it. This method is called importance sampling (Robert and Casella, 2013). In particular, here we can use the unit EPSP distribution *q*_*v*_ (*v*) as the proposal distribution, because unit EPSPs {*v*_*k*_} of synaptic connections can be interpreted as samples depicted from the unit EPSP distribution *q_v_* (Fig. 1B top). Thus, the mean 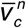is approximately calculated as

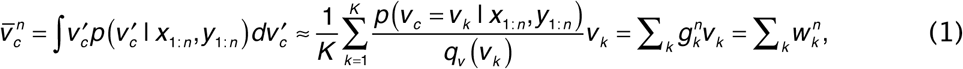

where 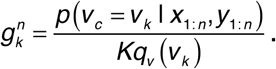 Therefore, if spine size*g*_*k*_^*n*^ represents the relative weight ofsample *v*_*k*_, then dendritic summation over postsynaptic potentials 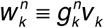 naturally represents the desired value 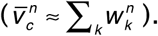 For instance, if the distribution of synapses is biased toward proximal side (i.e. if the mean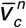 is overestimated by the distribution of unit EPSPs as in Fig. 1B top), then synapses at distal dendrites should possess large spine sizes, while the spine sizes of proximal synapses should be smaller (Fig. 1B bottom).

### Synaptic plasticity as particle filtering

In the previous section, we showed that redundant synaptic connections can represent probabilistic distribution *p*(*v*_*c*_=*v*_*k*_|*x*_*1:n*_,*y*_*1:n*_) if spine sizes {*g*_*k*_} coincide w6ith their importance 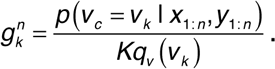 But, how can synapses update their representation of the probabilistic distribution *p*(*v*_*c*_=*v*_*k*_|*x*_*1:n*_,*y*_*1:n*_) based on a new observation {*x*_*n+1*_, *y*_*n+1*_}? Because *p*(*v*_*c*_=*v*_*k*_|*x*_*1:n*_,*y*_*1:n*_) is mapped onto a set of spine sizes {*g*_*k*_^*n*^} as in Equation 1, the update of the estimated distribution *p*(*v*_*k*_|*x*_*1:n*_,*y*_*1:n*_)→*p*(*v*_*k*_|*x*_*1:n+1*_,*y*_*1:n+1*_) can be performed by the update of spine sizes 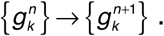. By considering particle filtering (Doucet et al., 2000) on the parameter space (see *The learning rule for multisynaptic connections* in Methods for details), we can derive the learning rule for spine size as

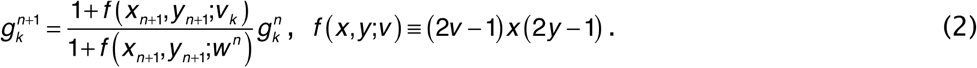

This rule is primary Hebbian, because the weight change depends on the product of pre-and postsynaptic activity *x*_*n+1*_ and *y*_*n+1*_. In addition to that, the change also depends on unit EPSP *v*_*k*_. This dependence on unit EPSP reflects the dendritic position dependence of synaptic plasticity. In particular, for a distal synapse (i.e. for small *v*_*k*_), the position-dependent term (2*v*_*k*_-1) takes a negative value (note that 0≤*v*_*k*_<1), thus yielding an anti-Hebbian rule as observed in neocortical synapses (Letzkus et al., 2006; Sjöström and Häusser, 2006).

For instance, if the new data {*x_n+1_*, *y_n+1_*} indicates that the value of *v*_*c*_ is in fact larger then previously estimated, then the distribution *p*(*v*_*c*_|*x*_*1:n+1*_,*y*_*1:n+1*_) shifts to the right side (upper panel of Fig. 1C). This means that the spine size *g*_*k*_^*n+1*^ becomes larger then *g_k_^n^* at synapses on the right side (i.e. proximal side), whereas synapses get smaller on the left side (i.e. distal side; bottom panel of Fig. 1C). Therefore, pre and postsynaptic activity causes LTP at proximal synapses induces LTD at distal synapses as observed in experiments (Letzkus et al., 2006; Sjöström and Häusser, 2006). The derived learning rule (Eq. 2) also depends on the total EPSP amplitude 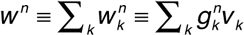. This term reflects a normalization factor possibly modulated through redistribution of synaptic vesicles over the presynaptic axon (Staras et al., 2010). A surrogate learning rule without this normalization factor will be studied in a later section.

We performed simulations by assuming that the two neurons are connected with ten synapses with the uniform unit-EPSP distribution (i.e. *q*_*v*_(*v*) = const.). At an initial phase of learning, the distribution of spine size {*g*_*k*_^*n*^} has a broad shape (purple lines in Fig. 1D), and the mean of distribution is far away from the true value (*v*=*v*_*c*_). However, the distribution is skewed around the true value as evidence is accumulated through stochastic pre and postsynaptic activities (red lines in Fig. 1D). Indeed, the estimation performance of the proposed method is nearly the same as that of the exact optimal estimation, and much better than the standard monosynaptic learning rules (Fig. 1E; see *Monosynaptic learning rule* in Methods for details).

### Synaptogenesis as resampling

As shown above, weight modification in multisynaptic connections enables a near optimal learning. However, to represent the distribution accurately, many synaptic connections are required (gray line in Fig. 2B), while the number of synapses between a excitatory neuron pair is typically around five in the cortical microcircuits. Moreover, even if many synapses are allocated between presynaptic and postsynaptic neurons, if the unit EPSP distribution is highly biased, the estimation is poorly performed (gray line in Fig. 2C). We next show that this problem can be avoided by introducing synaptogenesis (Holtmaat and Svoboda, 2009) into the learning rule.

**Fig. 2.**
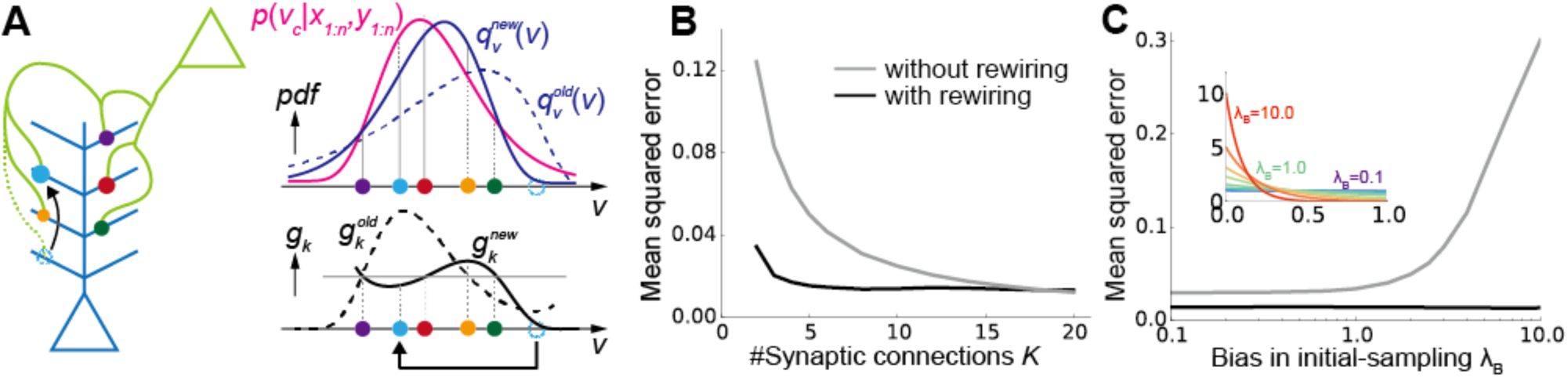
Synaptic rewiring for efficient learning. **A)** Schematic illustration of resampling. Dotted cyan circles represent an eliminated synapse, and the filled cyan circles represent a newly created synapse. **B, C)** Comparison of performance with/without synaptic rewiring at various synaptic multiplicity *K* **(B)**, and bias in initial-sampling *λ*_*B*_ **(C)**. For each bias parameter *λ*_B_, the unit EPSP distribution *{vk}* was set as 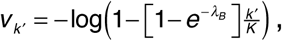 as depicted in the inset. Lines are the means over 104 simulations.

In the proposed framework, when synaptic connections are fixed (i.e. when *{vk}* are fixed), some synapses quickly become useless for representing the distribution. For instance, in Figure 2A, (dotted) cyan synapse is too proximal to contribute for the representation of *p*(*v*_*c*_|*x,y*). Therefore, by removing the cyan synapse and creating a new synapse at a random site, on average, the representation becomes more effective (Fig. 2A). Importantly, in our framework, spine size factor *gk* is proportional to the informatic importance of the synapse by definition, thus optimal rewiring is achievable simply by removing the synapse with the smallest spine size. Ideally, the new synapse should be sampled from *p*(*v*_*c*_|*x,y*) for an efficient rewiring, yet it is not clear if such a sampling is biologically plausible, and indeed random resampling is sufficient as long as elimination is selectively performed as mentioned above.

By introducing this resampling process, the model is able to achieve high performance even if the total number of synaptic connection is just around three (black line in Fig. 2B), or if the initial distribution of {*v*_*k*_} is poorly taken (black line in Fig. 2C).

### Detailed single neuron model of learning from many presynaptic neurons

In the previous sections, we found that synaptic plasticity in multisynaptic connections can achieve non-parametric near-optimal learning in a simple model with one presynaptic neuron. To investigate its biological plausibility, we next extend the proposed framework to a detailed single neuron model receiving inputs from many presynaptic neurons. To this end, we constructed an active dendritic model using NEURON simulator (Hines and Carnevale, 1997) based on a previous model of L2/3 pyramidal neurons of the primary visual cortex (Smith et al., 2013). We randomly distributed 1000 excitatory synaptic inputs from 200 presynaptic neurons on the dendritic tree of the postsynaptic neuron, while fixing synaptic connections per presynaptic neuron at *K*=5 (Fig. 3A; see *Morphology* in Methods for the details of the model). We assumed that all excitatory inputs are made on spines, and each spine is projected from only one bouton for simplicity. In addition, 200 inhibitory synaptic inputs were added on the dendrite to keep the excitatory/inhibitory (E/I) balance (Froemke, 2015). We first assigned a small constant conductance for each synapse, and then measured the somatic potential change, which corresponds to the unit EPSP in the model. As observed in cortical neurons (Stuart and Spruston, 1998), input at a more distal dendrite showed larger attenuation at the soma, though variability was quite high across branches (Fig. 3B).

**Fig. 3.**
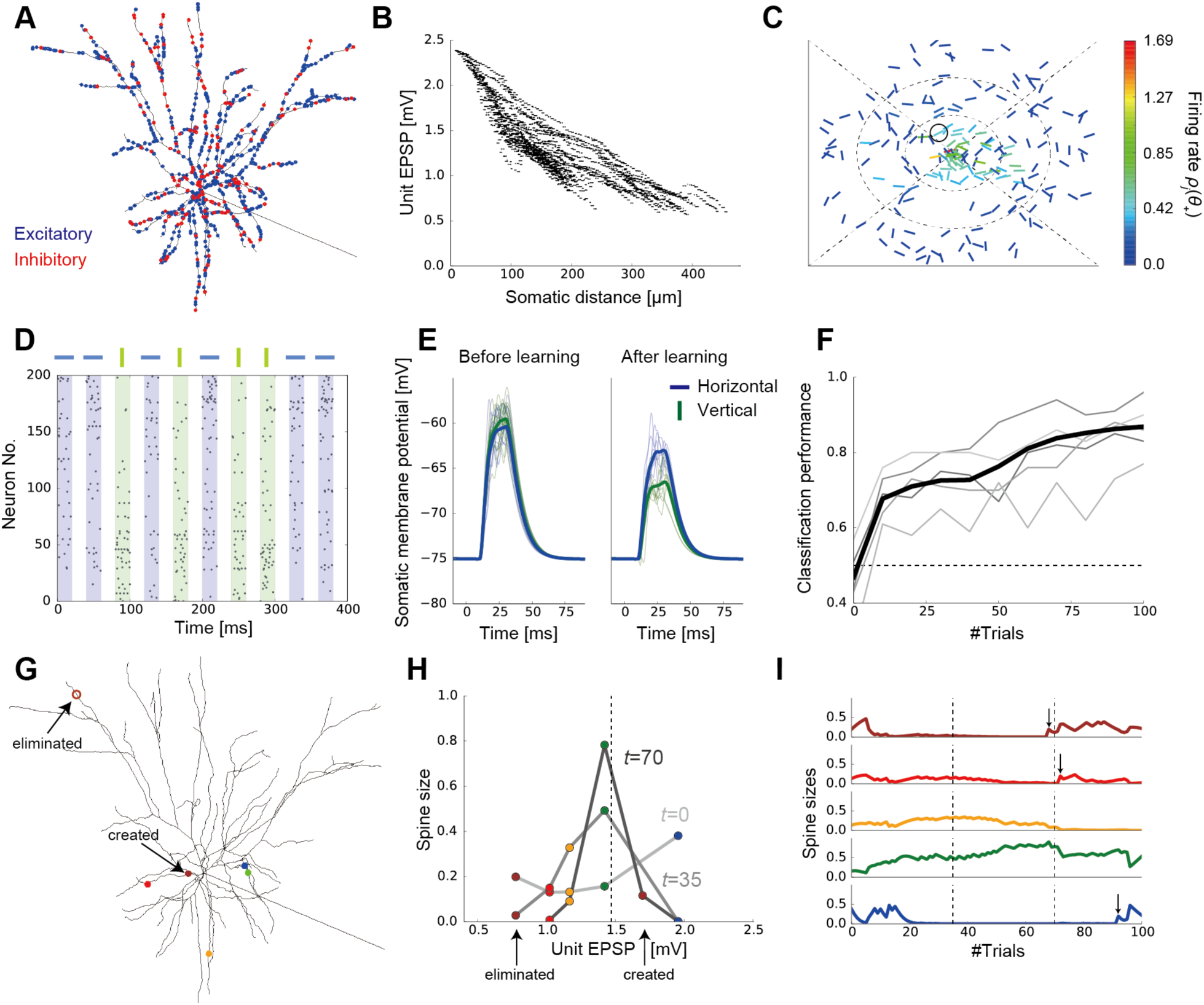
A detailed model of multisynaptic learning with multiple presynaptic neurons. **A)** Schematic figure of the detailed neuron model. Blue and red points on the dendritic trees represent excitatory and inhibitory synaptic inputs, respectively. **B)** Dendritic position dependence of unit EPSP. Each dot represents a synaptic contact on the dendritic tree. **C)** An example of the visual selectivity patterns of presynaptic neurons. Position and angle of each bar represent the receptive field (RF) and the orientation selectivity of each presynaptic neuron, where the RF was defined relative to the RF of the postsynaptic neuron (the central position). Colors represent the firing rates of presynaptic neurons when a horizontal bar stimulus is presented at the RF of the postsynaptic neuron. Here, the firing rates were evaluated as the expected number of spikes within 20ms stimulus duration (see *Stimulus selectivity* in Methods for details). The black circle shows the selectivity of the representative neuron depicted in **G-I. D)** Examples of input spike trains generated from the horizontal (target) and vertical (non-target) stimuli. Presynaptic neurons were sorted by their stimulus preference. Note that in the actual simulations, variables were initialized after each stimulation trial. See *Task configuration* in Methods for details of the task. **E** Somatic responses before and after learning. Thick lines represent the average response curves over 100 trials and thin lines are trial-by-trial responses. **F)** The average learning curves over 50 simulations (black line) and examples of learning curves (gray lines). **G-I)** An example of learning dynamics under the multisynaptic rule (see Results for details).

Next, we consider a perceptual learning task in this neuron model. Each excitatory presynaptic neuron was assumed to be a local excitatory neuron, modeled as a simple cell having a small receptive field (RF) and a preferred orientation in the visual space (Fig. 3C). Axonal projections from each presynaptic neuron were made onto five randomly selected dendritic branches of the postsynaptic neuron regardless of the stimulus selectivity, because visual cortex of mice has a rather diverse retinotopic structure (Bonin et al., 2011). In this setting, the post-neuron should be able to infer the orientation of the stimulus presented at its RF from the presynaptic inputs, because cells having similar RFs or orientation selectivity are often co-activated (Simoncelli and Olshausen 2001; Geisler et al., 2001). Thus, we consider a supervised learning task in which the postsynaptic neuron has to learn to detect a horizontal grading, not a vertical grading, from stochastic presynaptic spikes depicted in Figure 3D. In reality, the modulation of lateral connections in L2/3 is arguably guided by the feedforward inputs from layer 4 (Ko et al., 2013; Urbanczik and Senn 2014). However, for simplicity, we instead introduced an explicit supervised signal to the postsynaptic neuron. In this formulation, we can directly apply the rule for synaptic plasticity and rewiring introduced in the previous section (see *The learning rule for the detailed model* in Methods). Here, in addition to the rewiring by the proposed multisynaptic rule, we implemented elimination of synapses from uncorrelated presynaptic neurons, to better replicate developmental synaptic dynamics.

Initially, the postsynaptic somatic membrane potential responded similarly to both horizontal and vertical stimuli, but the neuron gradually learned to show a selective response to the horizontal stimulus (Fig. 3E). After 100 trials, the two stimuli became easily distinguishable by the somatic membrane dynamics (Fig. 3E and F; see *Performance evaluation* in Methods for details). Next, we examined how the proposed mechanism works in detail. To this end, we focused on a presynaptic neuron circled in Figure 3C, and tracked the changes in its synaptic projections and spine sizes (Fig. 3G-I). Because the neuron has a RF near the postsynaptic RF, and its orientation selectivity is nearly horizontal, the total synaptic weight from this neuron should be moderately large after learning. Indeed, the Bayesian optimal weight was estimated to be around 1.5 mV in the model (vertical dotted line in Fig. 3H), under the assumption of linear dendritic integration. Overall, the unit EPSPs of the majority of synapses were initially around 1.0-1.5 mV, while smaller or larger unit EPSPs were rare due to dendritic morphology (Fig. 3B). To counterbalance this bias toward the center, we initialized the spine size in a U-shape (light gray line in Fig. 3H). In this way, the prior distribution of the total synaptic weight becomes roughly uniform (see also Fig. 1B). After a short training, the most proximal spine (the blue one) was depotentiated, whereas spines with moderate unit EPSP sizes were potentiated (yellow and green ones on dark gray line in Fig. 3H). This is because, the expected distribution of the weight from this presynaptic neuron shifted to the left side (i.e. to a smaller EPSP) after the training, and this shift was implemented by reducing the spine size of the proximal synapse, while increasing the sizes of others (as in Fig. 1C, but here the change is to the opposite direction). Note that, the most distal spine (the brown one) was also depressed here, as the expected distribution got squeezed toward the center. Finally, after a longer training, the expected distribution became more squeezed, hence all but the green spine were depotentiated (black line in Fig. 3H). Moreover, the most distal synapse was eliminated because its spine size became too small to make any meaningful contribution to the representation, and a new synapse was created at a proximal site (open and closed brown circles in Fig. 3G, respectively) as explained in Figure 2A. This rewiring achieve a more efficient representation of the weight distribution on average. Indeed, the new brown synapse was potentiated subsequently (top panel in Fig. 3I). Note that, in this example, red and blue synapses were also rewired shortly after this moment (vertical arrows above red and blue traces in Fig. 3I).

### The model reproduces various properties of synaptic organization on the dendrite

While we confirmed that the proposed learning paradigm works well in a realistic model setting, we further investigated its consistency with experimental results. We first calculated spine survival ratio for connections from different presynaptic neurons. As suggested from experimental studies (Ko et al., 2013; Iacaruso et al., 2017), more synapses survived if the presynaptic neuron had a RF near the postsynaptic RF after learning (Fig. 4A). Likewise, synapses having similar orientation selectivity to the postsynaptic neuron showed higher survival rates (Fig. 4B) as indicated from previous observations (Ko et al., 2013; Lee et al., 2016). However, this orientation dependence was evident only for projections from neurons with a RF in the direction of the postsynaptic orientation selectivity (blue line in Fig. 4C), and the spines projected from neurons with orthogonal RFs remained to have uniform selectivity even after learning (green line in Fig. 4C), as reported in a recent experiment (Iacaruso et al., 2017). In contrast, both connections from neurons with nearby and faraway RFs showed clear orientation dependence, though the dependence was more evident for the latter in the model (Fig. 4D). The consistencies with the experimental results (Fig. 4A-D) support the legitimacy of our model setting, though they were achieved by the elimination of uncorrelated spines, not by the multisynaptic learning rule per se.

**Fig. 4.**
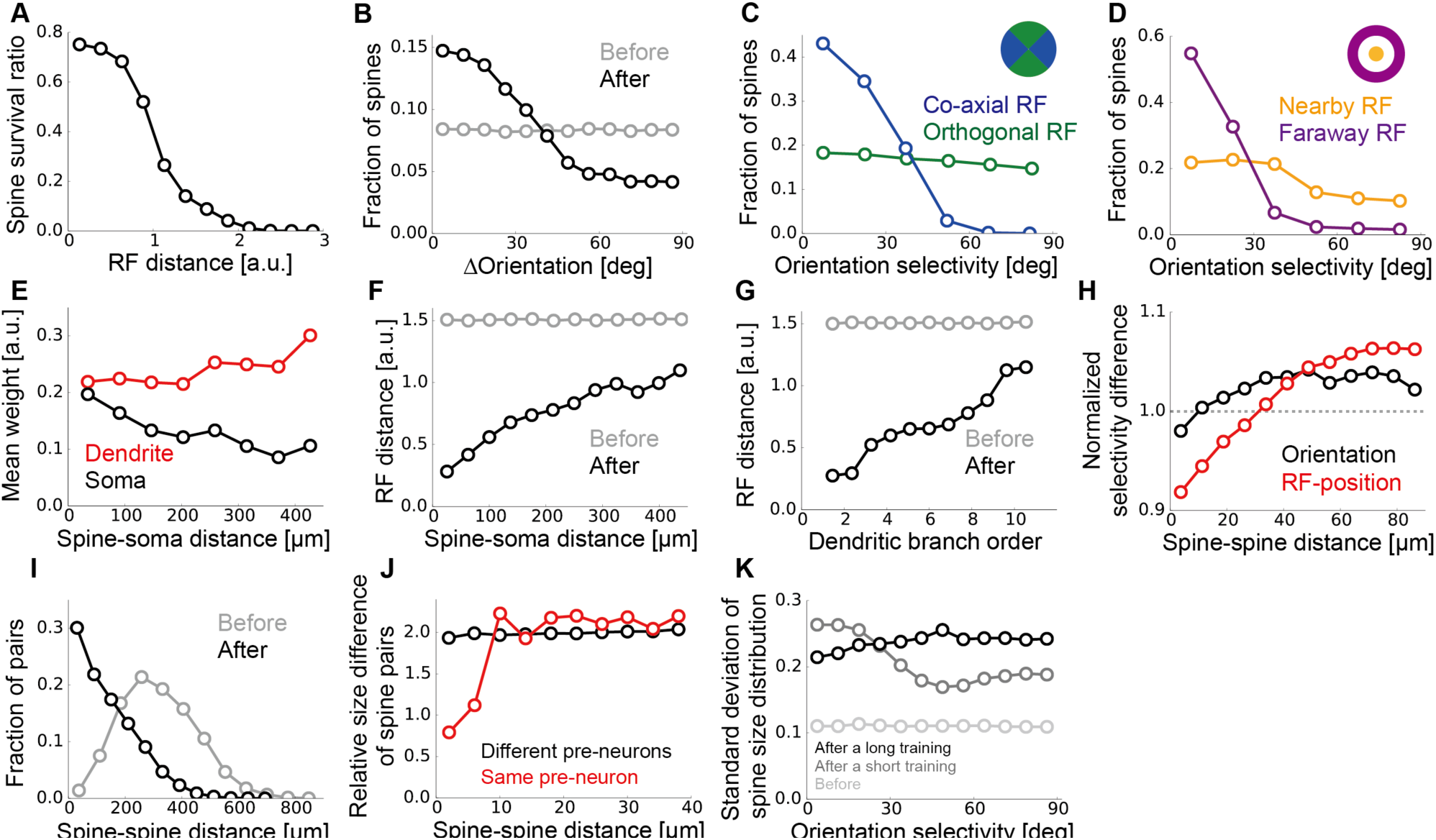
Synaptic organization on the dendrite by the multisynaptic learning rule. **A)** Survival ratio of spines with different receptive field (RF) distances from the postsynaptic neuron. **B)** Fraction of spines having various orientation selectivity before and after learning. **C, D)** Fraction of spines survived after learning, calculated for different orientation selectivity at co-axial/orthogonal RFs **(C)**, and at nearby/faraway RFs **(D)**. We defined the RF of presynaptic neuron *j* being orthogonal if 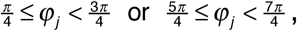 and co-axial otherwise. The RF of neuron *j* was defined as nearby if *r*_*j*_<0.5, but faraway if *r*_*j*_>1.0 (see *Stimulus selectivity* in Methods). **E)** Relationship between the dendritic distance and the relative weight at the dendrite *g*_*k*_ and the soma *g*_*k*_*v*_*k*_/*v*_*max*_. **F)** Relationship between the dendritic distance of a spine and its RF distance in the visual space. **G)** The same as **F**, but calculated for the dendritic branch order, not the dendritic distance. **H)** Dependence of normalized RF difference (red), and normalized orientation difference (black) on the between-spine distance were calculated for two synapses projected from different neurons. We used the Euclidean distance in the visual field 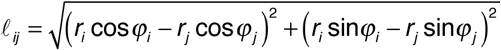for RF distance between presynaptic neurons *i* and *j*, and the normalization was taken over all synapse pairs. **I)** Distributions of dendritic distance between synapses projected from the same presynaptic neuron before and after learning. **J)** Relative spine size difference between spines projected from the same presynaptic neuron or different neurons calculated for pairs with different spine distance. The relative size difference between spine *i* and *j* was defined as |log(*gi/gj*)|. **K)** Standard deviation (SD) of spine size distribution at various orientation selectivity for synapses from presynaptic neurons with nearby RFs (*r*_*j*_<0.5). The distributions for short and long training were taken after learning from 10 and 1000 samples, respectively. All panels were calculated by taking averages over 500 independently simulated neurons, and the learning was performed from 1000 training samples.

We next investigated changes in dendritic synaptic organization generated by the multisynaptic learning. Overall, the mean spine size was slightly larger at distal dendrites (red line in Fig. 4E), but this trend was not strong enough to compensate the dendritic attenuation (black line in Fig. 4E), being consistent with previous observations in neocortical pyramidal neurons (Williams and Stuart, 2003). Importantly, neurons with RFs faraway from the postsynaptic RF likely formed synaptic projections more on distal dendrites than on proximal ones (Fig. 4F), and at higher dendritic branch orders than at lower ones (Fig. 4G), as observed previously (Iacaruso et al., 2017). This is because, in the proposed learning rule, if pre and postsynaptic neurons have similar spatial selectivity, synaptic connections are preferably rewired toward proximal positions (Fig. 3G), and vice versa (Fig. 2A). Moreover, nearby spines on the dendrite showed similar RF selectivity even if multisynaptic pairs (i.e., synapse pairs projected from the same neuron) were excluded from the analysis (red line in Fig. 4H), due to the dendritic position dependence of presynaptic RFs. On the other hand, similarity between nearby spines was less significant in orientation selectivity (black line in Fig. 4H), as observed previously in rodent experiments (Jia et al., 2010; Iacaruso et al., 2017). These results suggest a potential importance of developmental plasticity in somatic-distance dependent synaptic organization.

In the model, the position of a newly created synapse was limited to the branches where the presynaptic neuron initially had a projection, to roughly reproduce the spatial constraint on synaptic contacts. As a result, although there are many locations on the dendrite where the unit EPSP size is optimal for a given presynaptic neuron, only few of them are accessible from the neuron, hence synapses from the same presynaptic neuron may form clusters there. Indeed, by examining changes in multisynaptic connection structure, we found that the dendritic distance between two spines projected from the same presynaptic neuron became much shorter after learning (Fig. 4I), creating clusters of synapses from the same axon. This result suggests that clustering of multisynaptic connections observed in the experiments (Schmidt 2017) is possibly caused by developmental synaptogenesis under a spatial constraint. Furthermore, as observed in hippocampal neurons (Bartol et al., 2015), two synapses from the same presynaptic neuron had similar spine sizes if the connections were spatially close to each other, but the correlation in spine size disappeared if they were distant (red line in Fig. 4J). On the other hand, spine sizes of two synapses from different neurons were always uncorrelated regardless of the spine distance (black line in Fig. 4J).

Lastly, we studied the spine size distribution. In the proposed framework, the mean spine size does not essentially depend on presynaptic stimulus selectivity due to normalization, but the variance may change. In particular, the spine size variance is expected to be small if the presynaptic activity is highly stochastic, because the distribution of spine sizes stays nearly uniform in this condition, while the spine size variance should increase upon accumulation of samples. Indeed, in the initial phase of learning, the variance of spine size went up for projections from neurons with horizontal orientation selectivity (gray line Fig. 4K), though the spine size variance from other presynaptic neurons caught up eventually (black line Fig. 4K). In this regard, a recent experimental study found higher variability in postsynaptic density (PSD) areas for projections from neurons sharing orientation preference with the postsynaptic cell, though the data was from adult, not from juvenile mice (Lee et al., 2016).

### The multisynaptic rule robustly enables fast learning

The correspondence with experiment observations discussed in the previous section supports the plausibility of our framework as a candidate mechanism of synaptic plasticity on the dendrites. Hence, we further studied the robustness of learning dynamics under the proposed multisynaptic rule. Below, we turn off the spine elimination mechanism that is not compensated by creation, as this process affects the learning dynamics.

In the proposed model, if the initial synaptic distribution on the dendrite *q*_*v*_(*v*) is close to the desired distribution *p*_*v*_(*v*), spine size modification is in principle unnecessary. In particular, the optimal EPSPs of most presynaptic neurons are small in our L2/3 model (Fig. 3C); hence most synaptic contacts should be placed on distal branches on average. Indeed, when the initial synaptic distribution was biased toward the distal side, improvement in classification performance became faster (black vs blue lines in Fig. 5A). This result suggests that the synaptic distribution on the postsynaptic dendrite may work as a prior distribution.

**Fig. 5.**
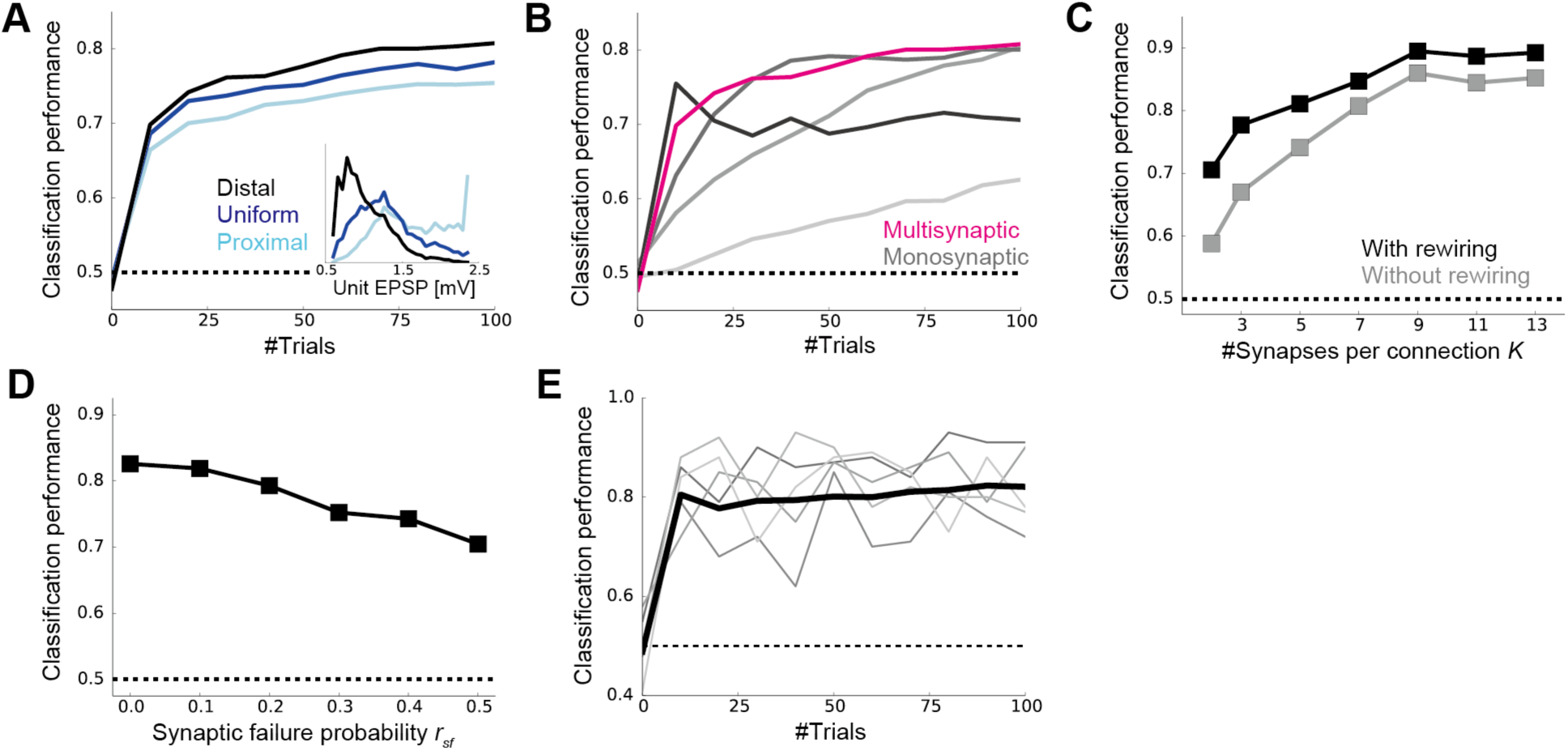
Dynamics of the multisynaptic learning rule under various conditions. **A)** Learning dynamics under various initial synaptic distributions. The inset represents the unit EPSP distributions when synaptic connections are biased toward the distal dendrite (black), unbiased (blue), and biased toward the proximal (light blue). **B)** Comparison with the monosynaptic learning. We set the learning rate as *η*_*w*_=0.03, 0.1, 0.3, 1.0, from light gray to black lines. To keep the E/I balance, the inhibitory weight was set to *γ*_*I*_=2.0 for *η*_*w*_=1.0, and *γ*_*I*_=1.25 for the rest. The magenta line is the same as the black line in **A. C)** Classification performance after learning with different numbers of synapses per connection with or without rewiring. For the E/I balance, the inhibitory weights were chosen as *γ_I_*=2.0, 1.2, 0.75, 0.6, 0.5, 0.4, 0.3, when the number of synapses per connections were *K*=2, 3, 5, 7, 9, 11, 13, respectively. **D)** The performance after learning with various synaptic failure probabilities. Both in panel **C** and **D**, the performance was calculated after 1000 trials. **E)** Learning dynamics under the surrogate rule. Thin gray lines represent examples. All panels were calculated by taking the means over 50 simulations.

We next compared the learning performance with the standard monosynaptic learning rule in which the learning rate is a free parameter (see *Monosynaptic rule for the detailed model* in Methods). If the learning rate is chosen at a small value, the neuron took a very large number of trials to learn the classification task (light gray line in Fig. 5B). On the other hand, if the learning rate is too large, the learning dynamics became unstable and the performance dropped off after a dozen trials (black line in Fig. 5B). Therefore, the learning performance was comparable with the multisynaptic rule only in a small parameter region (*η*_*w*_~0.1). By contrast, in the multisynaptic rule, stable fast learning was achievable without any fine-tuning (magenta line in Fig. 5B).

As expected from Figure 2, the proposed learning mechanism worked well even if the number of synapses per connection was small (Fig. 5C). Without rewiring, the classification task required seven synapses per connection for an 80% success rate, but three was enough with rewiring (Fig. 5C). Moreover, the learning performance was robust against synaptic failure (Fig. 5D). Although local excitatory inputs to L2/3 pyramidal cells have a relatively high release probability (Branco and Staras, 2009), the stochasticity of synaptic transmission at each synapse may affect learning and classification. We found that even if the half of presynaptic spikes were omitted at each synapse (see *Task configuration* in Methods for details), the classification performance was still significantly above the chance level (Fig. 5D).

In the proposed model, competition was assumed among synapses projected from the same presynaptic neuron, but it is unclear if homeostatic plasticity works in such a specific manner. Thus, we next constructed a surrogate learning rule that only requires a global homeostatic plasticity. In this rule, the importance of a synapse was not compared with other synapses from the same presynaptic neuron, but was compared with a hypothesized standard synapse (see *The surrogate learning rule* in Methods). When the unit EPSP size of the standard synapse was chosen appropriately, the surrogate rule indeed enabled neuron to learn the classification task robustly and quickly (Fig. 5E). Overall, these results support the robustness and biological plausibility of the proposed multisynaptic learning rule.

## Discussion

In this work, first we have used a simple conceptual model to show: (i) Multisynaptic connections provide a non-parametric representation of probabilistic distribution of the hidden parameter using redundancy in synaptic connections (Fig. 1AB); (ii) Updating of probabilistic distribution given new inputs can be performed by a Hebbian-type synaptic plasticity when the output activity is supervised (Fig. 1C-E); (iii) Elimination and creation of spines is crucial for efficient representation and fast learning (Fig. 2A-C). In short, synaptic plasticity and rewiring at multisynaptic connections naturally implements an efficient sample-based Bayesian filtering algorithm. Secondly, we have demonstrated that the proposed multisynaptic learning rule works well in a detailed single neuron model receiving stochastic spikes from many neurons (Fig. 3). Moreover, we found that the model reproduces the somatic-distance dependent synaptic organization observed in the L2/3 of rodent visual cortex (Fig. 4F and G). Furthermore, the model suggests that the dendritic distribution of multisynaptic inputs provides a prior distribution of the expected synaptic weight (Fig. 5A).

### Experimental predictions

Our study provides several experimentally testable predictions on dendritic synaptic plasticity, and the resultant synaptic distribution. First, the model suggests a crucial role of developmental synaptogenesis in the formulation of presynaptic selectivity-dependent synaptic organization on the dendritic tree (Fig. 4F and G), observed in the primary visual cortex (Iacaruso et al., 2017). More specifically, we have revealed that the RF-dependence of synaptic organization is a natural consequence of the Bayesian optimal learning under the given implementation. Evidently, retinotopic organization of presynaptic neurons is partially responsible for this dendritic projection pattern, as a neuron tends to make a projection onto a dendritic branch near the presynaptic cell body (Markram et al., 2015; Gal et al., 2017). However, a recent experiment reported that RF-dependent global synaptic organization on the dendrite is absent in the primary visual cortex of ferrets (Scholl et al., 2017). This result indirectly supports the non-anatomical origin of the dendritic synaptic organization, as a similar organization is arguably expected in ferrets if the synaptic organization is purely anatomical.

Our study also predicts developmental convergence of synaptic connections from each presynaptic neuron (Fig. 3G and Fig. 4I). It is indeed known that in adult cortex, synaptic connections from the same presynaptic neuron are often clustered (Kasthuri et al., 2015; Schmidt, 2017). Our model interprets synaptic clustering as a result of an experience-dependent resampling process by synaptic rewiring, and predicts that synaptic connections are less clustered in immature animal. In particular, our result suggests that synaptic clustering occurs in a relatively large spatial scale (~100μm; as shown in Fig 5I), not in a fine spatial scale (~10μm). This may explain a recent report on the lack of fine clustering structure in the rodent visual cortex (Lee et al., 2016).

Furthermore, our study provides an insight on the functional role of anti-Hebbian plasticity at distal synapses (Letzkus et al., 2006; Sjöström and Häusser, 2006). Even if the presynaptic activity is not tightly correlated with the postsynaptic activity, that does not mean the presynaptic input is not important. For instance, in our detailed neuron model, inputs from neurons having a RF faraway from the postsynaptic RF still helps the postsynaptic neuron to infer the presented stimulus (Fig. 3). More generally, long-range inputs are typically not correlated with the output spike trains, because the inputs usually carry contextual information (Bittner et al., 2015), or delayed feedback signals (Manita et al., 2015), yet play important moduratory roles. Our study indicates that anti-Hebbian plasticity at distal synapses prevents these connections from being eliminated, by keeping the synaptic connection strong. This may explain why modulatory inputs are often projected to distal dendrites (Bittner et al., 2015; Manita et al., 2015), though active dendritic computation shuold also be crucial especially in case of Layer 5 or CA1 pryramidal neurons (Segev and London, 2000).

### Related works

Previous theoretical studies often explain synaptic plasticity as stochastic gradient descent on some objective functions (Pfister et al., 2006; Nessler et al., 2013; Urbanczik and Senn, 2014; Hiratani and Fukai, 2016), but these models require fine-tuning of the learning rate for explaining near-optimal learning performance observed in humans (Behrens et al., 2007; Lake et al., 2015) and rats (Madarasz et al., 2016), unlike our model. Moreover, in this study, we proposed synaptic dynamics during learning as a sample-based inference process, in contrast to previous studies in which sample-based interpretations were applied for neural dynamics (Orbán et al., 2016).

On the anti-Hebbian plasticity at distal synapse, previous modeling studies have revealed its potential phenomenological origins (Graupner and Brunel, 2012), but its functional benefits, especially optimality, have not been well investigated before. Particle filtering is an established method in machine learning (Doucet et al., 2000), and has been applied to artificial neural networks (Freitas et al., 2000), yet its biological correspondence had been elusive.

Previous computational studies on dendritic computation have been emphasizing the importance of active dendritic process (Segev and London, 2000), especially for performing inference from correlated inputs (Ujfalussy et al., 2015), or for computation at terminal tufts of cortical layer 5 or CA1 neurons (Urbanczik and Senn, 2014). Nevertheless, experimental studies suggest the summation of excitatory inputs through dendritic tree is approximately linear (Cash and Yuste, 1999; Hao et al., 2009). Indeed, we have shown that a linear summation of synaptic inputs is suitable for implementing importance sampling. Moreover, we have demonstrated that even in a detailed neuron model with active dendrites, a learning rule assuming a linear synaptic summation works well.

## Methods

### A conceptual model of multisynaptic learning

#### The learning rule for multisynaptic connections

In the model, CS (eg. tone stimulus) and US (eg. electric shock) were represented by binary variables *x*_*n*_*∈*{0,1} and *y*_*n*_ *∈*{0,1}. At each trial *n*, CS was delivered with Pr[*x*_*n*_ = 1] = *π*_*x*_, and US was given only when *x*_*n*_=1, with probability Pr[*y*_*n*_= 1|*x*_*n*_= 1] = *v*_*c*_. For this task, the update rule for the spine size factor 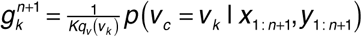 is given as,

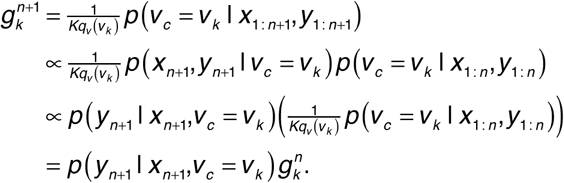

In particular, in our problem setting, *v*_*c*_ does not provide any information about *y*_*n*_ when *x*_*n*_=0, thus approximately (see the proof of convergence below),

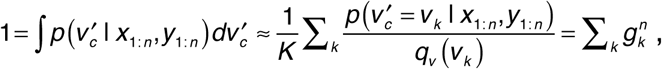

Because the normalization factor is determined by

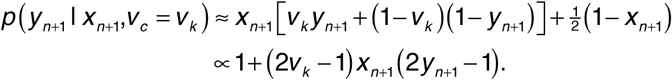

the sum of *{*g*_*k*_^*n*+1^}* should also be normalized to 1. Thus the update rule is given as

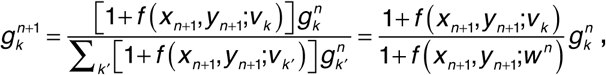

where *f*(*x,y*; *v*) ≡ (2*v*-1)*x*(2*y*-1) and 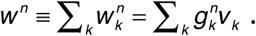 As for the resampling process, at every trial *n*, if spine *k* satisfied *g*_*k*_ < *g*_*th*_, unit EPSP was resampled uniformly from [0,1), and the spine size was set to *g*_*k*_ = *g*_*th.*_

#### Proof of convergence

The derived learning rule can be rewritten as

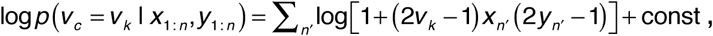

so in order to prove convergence, we need to show that 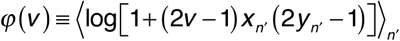 is maximized at true *v*_*c*_. By considering Taylor expansion, the above equation is expanded as 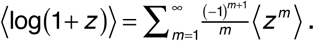In this form, the average is calculated as

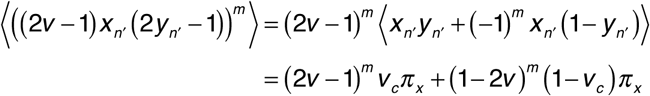

Note that (x_n_)^m^=x_n_ if m>0, because x_n_=0 or 1. Thus, by substituting the above equation into the Taylor expansion form,

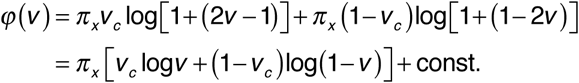

Therefore, φ(v) is maximized at *v* = *v*_*c*_.

#### Monosynaptic learning rule

For comparison, we implemented a monosynaptic learning rule. By expanding the exact solution 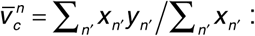

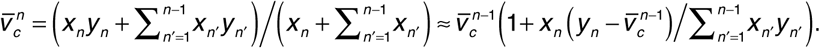

Hence, by using a single variable *v*_*m*_^*n*^, the learning rule is given as 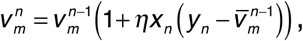
where *η* represents the learning rate. In the optimal learning depicted in Figure 1E, *v*_*c*_ was estimated as 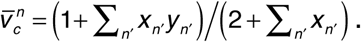

#### Details of the conceptual model

In the simulations, we used *π*_*x*_=0.3, and *v*_*c*_ was randomly chosen from [0,1) uniformly at each simulation (not at each trial). The number of connections was kept at *K*=10 except for Figure 2B in which *K*=2 to 20 were used. Initial value of *k-th* connection *v*_*k*_ was set as *v*_*k*_=(*k*+0.5)/*K* except for Figure 2C in which the initial distribution was biased by choosing 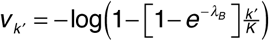 where ***\**B* is the bias parameter. Resampling was performed with the threshold *g_th_*=0.0001, and a new unit EPSP *v*_*k*_ was uniformly sampled from [0,1). In Figure 2B and C, the errors were calculated after learning from 10^4^ trials.

## Detailed single neuron model

### Morphology

We constructed a detailed neuron model based on a model of L2/3 pyramidal neuron with active dendrites (Smith et al., 2013) using NEURON simulator (Hines and Carnevale, 1997). Here, we used the original reconstructed morphology without scaling. We distributed 1000 excitatory synaptic inputs from 200 presynaptic neurons randomly on the dendrite. Synaptic input was modeled as a double exponential conductance change with the rise time *τ*_*rise*_=0.5ms, the decay time *τ*_*decay*_=2.5ms, and the reversal potential was set to 0mV. For each synapse *k* from presynaptic neuron *j*, we first applied a synaptic input with a constant weight factor *γ*_*g*_=2.5nS, and then determined the unit EPSP 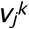 of synapse *k* by measuring somatic membrane potential change. The minimum and the maximum value of the unit EPSP of the given model were *v*_*min*_=0.57mV and *v*_*max*_=2.39mV, respectively. In the simulation of the task, using malleable spine size factor 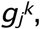 we set the weight factor of synapse *k* as 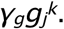 Similarly, 200 inhibitory synaptic inputs were uniformly distributed on the dendrite, and the rise and decay time of conductance was set as 0.5ms and 2.5ms, and the reversal potential was set to −90mV. The inhibitory weight factor was chosen as *γ*_*I*_=0.75nS.

### Stimulus Selectivity

We hypothesized that all excitatory presynaptic neurons are simple cells having direction selectivity{*θ*_*j*_} at receptive field (RF) {(*θ*_*j*_, *φ*_*j*_)}. Here, the position of RF in the visual field was defined by the relative position to the postsynaptic neuron in the polar coordinate (Fig. 3C). We modeled the mean firing rate of presynaptic neuron *j* for a stimulus *θ* at the RF of the postsynaptic neuron (i.e. at *r*=0) as

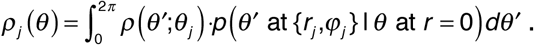

The first term *ρ*(*θ*’;*θ*_*j*_) is the mean response of the neuron with orientation selectivity *θ*_*j*_ when orientation *θ*’ is presented at its own RF, hence using a von Mises distribution, the response is approximately given as 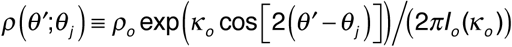(Swindale,1998). The second term is the probability of observing a stimulus with orientation *θ*’ at the position (*r*_*j*_, *φ*_*j*_) given stimulus *θ* at the center. The orientation *θ*’ at (*r*_*j*_, *φ*_*j*_) should be similar to the orientation *θ* at the center if *r*_*j*_ ~ 0, or *φ*_*j*_ ~ *θ* due to continuity and contour statistics (Simoncelli and Olshausen 2001; Geisler et al., 2001). Hence, we modeled the conditional probability as

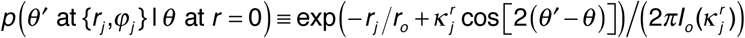

where 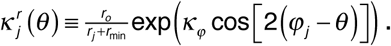 Note that the marginalized probability exp(-*r*_*i*_/*r*_*o*_) is smaller than one as an explicit stimulus may not exist at (*r*_*j*_, *φ*_*j*_) if the RF is far away from the center. By calculating the integral, the mean firing rate is derived as 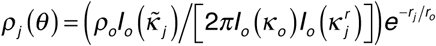 where 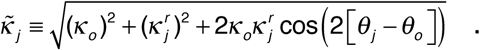. In the simulation, we used *κ*_*o*_=2.0, *κφ* =4.0, *ρ*_*o*_=1.5n, *r*_*min*_ =0.01exp(*κφ*), and *ρ*_*o*_ =1.0. The selectivity of each presynaptic neuron was uniformly sampled from the ranges: 0≤*r*_*j*_<3, *0≤φ*_*j*_<2*π*, and 0 ≤*θ*_*j*_<*π*.

Based on the selectivity described above, we modeled the spiking activity of presynaptic neuron *j* as a Poisson process with the rate *ρ*=*ρ*_*j*_(*θ*) under the presence of stimulus *θ*=*θ+* or *θ-.* In addition, we assumed that all presynaptic neurons follow a Poisson process with the rate *ρ*=*ρ*_*sp*_ in the spontaneous activity. In the simulation, we set *ρ*_*sp*_=0.01*ρ*_*o*_.

### Task configuration

We next consider the activity of the postsynaptic neuron. A sensory neuron should decode the presented stimulus given stochastic spiking spikes of presynaptic neurons. In particular, here we consider decoding of stimulus orientation *0* given spike counts from *M* presynaptic neurons *s*_*1*_:*M*^*t*^={*s*_*1*_^*t*^,*s*_*2*_^*t*^,…,*s*_*M*_^*t*^}. As the spikes were generated from Poisson processes in the model, the log-likelihood ratio of *θ*=*θ+* against the spontaneous activity *ϕ* is given as

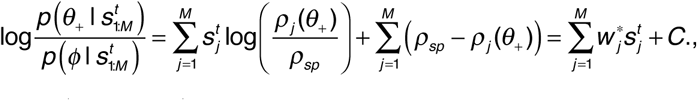

where 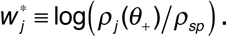 Hence, if the synapses projected from presynaptic neuron *j* learn to represent *w*_*j*_^*^ jointly, the somatic membrane potential naturally represents the log-likelihood of the stimulus being *θ*+, assuming passive dendritic integration.

In this task configuration, the estimated log-likelihoods are on average the same for two perpendicular stimuli *θ*=*θ*+ and *θ*- before learning, but the estimated log-likelihood becomes significantly larger for *θ*=*θ*+ once the correct weight structure is acquired. Hence, we evaluated the learning performance by a classification between *θ*=*θ*+ and *θ*-, using *θ*- as a control.

In the simulation, we first generated the spike counts of each presynaptic neurons {*s*_*1*_^*t*^, *s*_*2*_^*t*^,…,*s*_*M*_^*t*^} by sampling from Poisson distributions with the rates {*ρ*_*1*_, *ρ*_*2*_,…, *ρ*_*M*_} where *ρ*_*j*_=*ρ*_*j*_ (*θ+*) or *ρ*_*j*_(*θ-*) depending on the task. Based on the spike count *s*_*j*^*t*^_, spike timings of the *m*-th spike from presynaptic neuron *j* at trial *t* was determined as 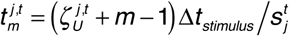where *Δt*_*stimulus*_=20ms, and 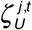 is a random variable uniformly depicted from [0,1). In the presence of synaptic failure, we instead defined a spike count at each synapse *k* by 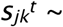 Binomial(*s*_*j*^*t*^_, 1-*r*_*sf*_), where *r*_*sf*_ is the failure rate. Inhibitory spikes were calculated in the same way, but the spike probability was defined by the total excitatory inputs as 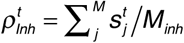 to achieve the E/I balance.

### The learning rule for the detailed model

We next derived the multisynaptic learning rule for this task. The optimal estimation of the weight from presynaptic neuron *j* at trial *t* is given as

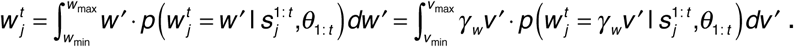

Here, we introduced a scaling factor *γ*_*w*_ to represent a dimensionless value *w* by a unit EPSP *v* [mV]. In the simulation, we used *γ*_*w*_=w_max/v__max_. By importance sampling,

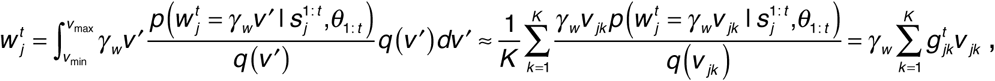

where 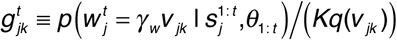represents the relative spine size of spine *k* from presynaptic neuron *j*, and *K* is the total number of synapses per presynaptic neuron. Therefore, considering a Bayesian filtering, the update of {*w*^*t*^_*j*_} is done by the following update of spine size {*g*^*t*^_*jk*_}

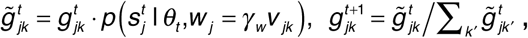

where 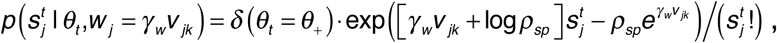 and *δ*(*x*) is a function that returns 1 if *x* is true, but returns 0 otherwise.

At every trial, synapses with spine size *g*_*jk*_^*t*^ < *g*_*th*_ was removed with 20% chance. If a synapse is removed, a new synaptic contact from the corresponding presynaptic neuron was simultaneously created on one of the dendritic branches to which the neuron initially had projections. Probability of selecting a branch was set to be proportional to the length of the branch. Spine size of a newly created synapse was set to *g*_*jk*_^*t*^=1/K. This rewiring procedure is slightly different from the one in the conceptual model, because rewiring becomes too frequent if we directly apply the latter.

In addition to rewiring of synaptic connections, we also included an elimination process that is not compensated by new connections, as the total number of synaptic connections is known to decreases during development (Holtmaat and Svoboda, 2009). In particular, inactive synapses are expected to be more fragile (Wiegert and Oertner, 2013). Hence, we tracked the firing rate of presynaptic neuron during the training phase by 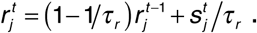. At every trial, if the presynaptic firing rate satisfies *r*_*j*_^*t*^ < *r*_*el-th*_, we eliminated the synaptic contact with 20% chance. Throughout the simulation, we used *g*_*th*_ = 0.001, *τr*=10.0, and *r*_*el-th*_=0.05.

### Monosynaptic learning rule for the detailed model

As presynaptic neurons follow stationary Poisson processes, the learning rule for monosynaptic connection was defined as

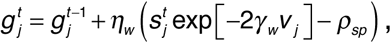

where *η*_*w*_ is the learning rate parameter (Nessler et al., 2013; Hiratani and Fukai, 2016), and *v*_*j*_ is the unit EPSP of the synaptic connection from neuron *j*. To ensure stability, we bounded the spine size between 0 < *g*_*j*_^*t*^ 0<1, and doubled the scaling factor from *γ*_*w*_ to 2*γ*_*w*_.

### The surrogate learning rule

In the surrogate rule, each synapse estimates the mean unit EPSP by 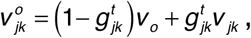 where *v*_*o*_ is thestandard unit EPSP. Subsequently, a synapse updates its spine size by

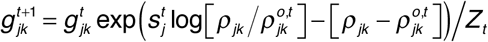

where 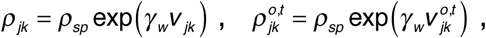, and 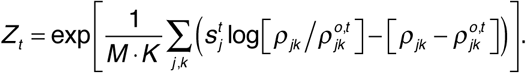 The normalization term *Z_t_* is global in a sense that the term is given by the summation over all the excitatory synapses projected to the postsynaptic neuron. To ensure the stability, we bounded the spine size factor as 0≤*g*^*t*^_*jk*_≤1/2,and set *v*_*o*_=1.5v_min_ (≈0.9mV).

### Performance evaluation

During the training phase, only the target (i.e. horizontal stimulus *θ*=*θ+*) was presented. In the test phase, we presented 200 stimuli, of which 100 stimuli were the horizontal stimulus (*θ*=*θ+*), while the other half were the vertical stimulus (*θ*=*θ-*). In Figure 3F, 5A, 5B and 5E, we stopped the training at every 10 trials, and measured the performance. The classification performance was measured by the ratio of horizontal trials in which the maximum EPSP height 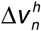 exceeded the threshold 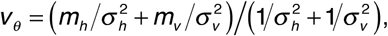 to the total of 100 trials, where 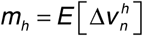 and 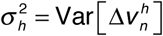 were calculated over 100 test stimuli (*n*=1, 2,…,100). Although the evaluations were made solely on false negatives, we also observed significant decrease of false positives during learning (Fig. 3E). When a postsynaptic action potential was emitted, we used the estimated membrane threshold *Δv*_*th*_=25mV as the maximum EPSP height Δ*v*_*n*_, but such a trial was rare (<1%) in our model setting.

### Details of the NEURON simulations

Initial values of spine sizes {*g*_*j*_^*k*^} were chosen such that 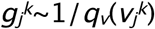 is satisfied. To this end, we first estimated the unit EPSP density at *v*=*v*_*j*_^*k*^ through a sample-based approximation:

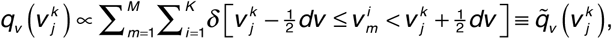

where *dv*=(*v*_*max*_-*v*_*min*_)/10. Then we calculated 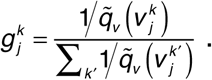 In Figure 5A, to generate a biased synaptic distribution, we randomly sampled a position from the whole dendritic tree with probability 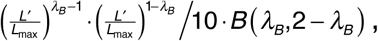 and added a synapse until 1000 synapses are created on the dendritic tree. Here, *L’* is the distance from the soma, *L*_*max*_ is its maximum length, *λ*_*B*_ is the bias parameter, and B(x,y) is the Beta function.

Presynaptic selectivity and initial synaptic contacts were randomly generated for each simulation, while the dendritic morphology was fixed. Further details of the model are available at ModelDB (modeldb.yale.edu/225075 with access code “1234”).

## Acknowledgements

The authors thank to Peter Latham for discussions and comments on the manuscript. This work was partly supported by CREST, JST (JPMJCR13W1 to TF) and Grants-in-Aid for Scientific Research (KAKENHI) from MEXT (no 15H04265 and 16H01289 to TF).

